# Woodsmoke particulates alter expression of antiviral host response genes in human nasal epithelial cells infected with SARS-CoV-2 in a sex-dependent manner

**DOI:** 10.1101/2021.08.23.457411

**Authors:** Stephanie A. Brocke, Grant T. Billings, Sharon Taft-Benz, Neil E. Alexis, Mark T. Heise, Ilona Jaspers

## Abstract

We have previously shown that exposure to particulate air pollution, both from natural and anthropogenic sources, alters gene expression in the airways and increases susceptibility to respiratory viral infection. Additionally, we have shown that woodsmoke particulates (WSP) affect responses to influenza in a sex-dependent manner. In the present study, we used human nasal epithelial cells (hNECs) from both sexes to investigate how particulate exposure could modulate gene expression in the context of SARS-CoV-2 infection. We used diesel exhaust particulate (DEP) as well as WSP derived from eucalyptus or red oak wood. HNECs were exposed to particulates at a concentration of 22 μg/cm^2^ for 2 h then immediately infected with SARS-CoV-2 at a MOI (multiplicity of infection) of 0.5. Exposure to particulates had no significant effects on viral load recovered from infected cells. Without particulate exposure, hNECs from both sexes displayed a robust upregulation of antiviral host response genes, though the response was greater in males. However, WSP exposure before infection dampened expression of genes related to the antiviral host response by 72 h post infection. Specifically, red oak WSP downregulated *IFIT1, IFITM3, IFNB1, MX1, CCL3, CCL5, CXCL11, CXCL10*, and *DDX58*, among others. After sex stratification of these results, we found that exposure to WSP prior to SARS-CoV-2 infection downregulated anti-viral gene expression in hNECs from females more so than males. These data indicate that WSP, specifically from red oak, alter virus-induced gene expression in a sex-dependent manner and potentially suppress antiviral host defense responses following SARS-CoV-2 infection.

## Introduction

Several concurrent natural disasters occurred in 2020 in the United States and abroad. Record-breaking wildfires ravaged the western United States and the COVID-19 pandemic posed a substantial public health burden around the world. Over 10 million acres of land were burned by wildfires in the United States in 2020 (1). This marked the second-highest acreage burned in a single year since the National Interagency Fire Center began recording in 1983 (1). Indeed, the area burned each year by wildfires has been trending upward in the United States for several decades (2).

Wildfires contribute significantly to air pollution and ambient particulate matter (PM). Particulate air pollution released from burning wildlands is associated with negative respiratory and cardiovascular health outcomes (reviewed in (3–6)) the toxicity of which depends heavily on the type of biomass burned and the burn temperature (7). Wildland firefighters and people who live downwind of or near to wildfires are exposed to high levels of PM released by fires (8). Studies have shown that wildland firefighters can be exposed to respirable particulate matter at concentrations >1 mg/m^3^ over the course of their work shift with maximum exposures reaching >2.5 mg/m^3^ (9–11). Large, populous regions of the western United States were exposed to unhealthy (>150 μg/m^3^) or hazardous (>300 μg/m^3^) air quality from PM_2.5_ and PM_10_ during September of 2020 (Fig. 1) (12). Epidemiological studies examining the health effects of wildfires showed an association between PM_2.5_ from wildfires and increased respiratory hospitalizations across 16 western states (13). A similar health effects study in California showed that women were more likely than men to visit the hospital for asthma- or hypertension-related reasons due to an increase in wildfire-generated PM_2.5_ (14).

**Fig. 1:**
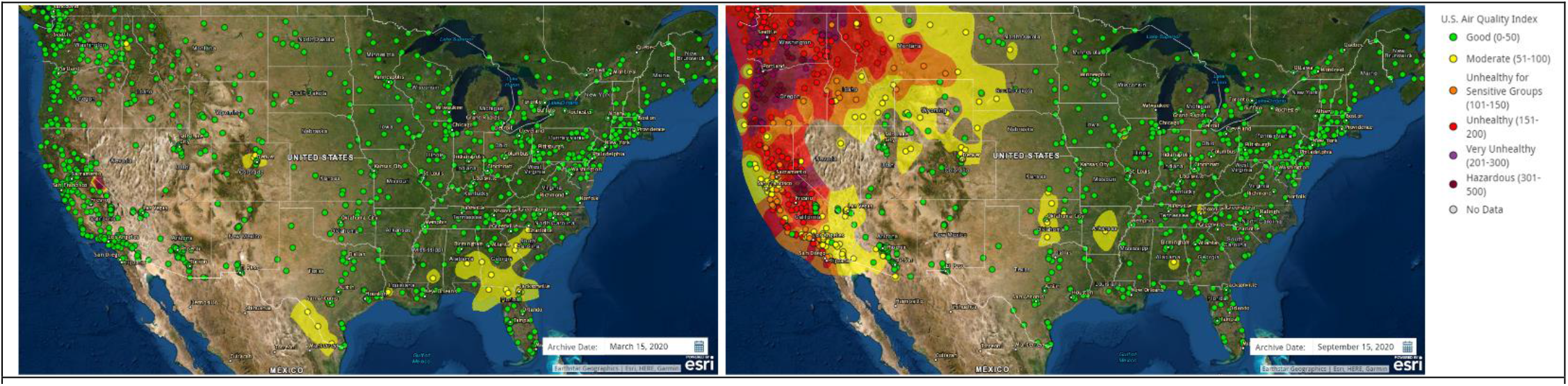
Air quality in the US on March 15, 2020 (left) and September 15, 2020 (right), before and during the 2020 fire season, respectively. Air quality monitoring stations (dots) and contours report the daily air quality index as defined by the US Environmental Protection Agency based on PM2.5 and PM10. Images retrieved from AirNow (https://gispub.epa.gov/airnow/index.html?tab=3).

Coinciding with record-breaking wildfires is the global coronavirus disease 2019 (COVID-19) pandemic that is, to date, responsible for over 3.8 million deaths worldwide (15). Sex has been found to affect COVID-19 outcomes with males more likely than females to develop severe or fatal cases of the disease (16–18).

SARS-CoV-2, the etiologic agent behind COVID-19, primarily affects the respiratory system (19) and exhibits tropism for cells of the upper airways, with nasal epithelial cells displaying the highest susceptibility to infection compared to bronchial and lower airway epithelium (20). Primary human nasal epithelial cells (hNECs) grown *in vitro* at air-liquid interface mimic *in vivo* differentiation patterns, evidenced by expression of mucins, presence of beating cilia, and tight junction formation (21, 22). Because the nasal epithelium expresses the SARS-CoV-2 viral entry factors ACE2 and TMPRSS2 in ciliated and secretory cells (23), the differentiated hNEC model is an excellent *in vitro* culture system to study SARS-CoV-2 pathogenesis. Along with biological aerosols, nasal epithelial cells are exposed to airborne particulates, gaseous pollution, and allergens. Thus, in addition to being a suitable model for studying respiratory viral infection (24, 25), hNECs demonstrate utility for toxicological studies involving aerosolized (26, 27), gaseous (28), and particulate (24) toxicants.

Exposure to air pollution is known to alter susceptibility to respiratory viral infection (reviewed in (29, 30)). *In vitro* models of respiratory epithelium treated with diesel exhaust particulate (DEP) prior to influenza infection increased viral attachment and the number of virus-infected cells relative to untreated cells (24). Rebuli and colleagues recently showed that red oak woodsmoke exposure followed by live attenuated influenza virus (LAIV) inoculation suppressed expression of host defense genes in women and upregulated many pro-inflammatory genes in men (31). Numerous epidemiological studies from around the world have found correlations between ambient air pollution levels and COVID-19 case number or case fatality rate (32–36). Recently, two studies found positive associations between ambient wildfire-derived PM_2.5_ and COVID-19 cases and deaths in the western United States (37, 38).

The present study examined the interactive effects of sex, exposure to WSP, and SARS-CoV-2 infection on gene expression in hNECs. To do this we exposed hNECs derived from healthy human donors to DEP or WSP derived from burned eucalyptus or red oak. Particulate exposures occurred prior to infection with SARS-CoV-2 and sampling occurred at 0, 24, and 72 h post infection (p.i.). We assembled a panel of 46 genes related to respiratory viral infection and host immune response, including the SARS-CoV-2 entry factor (ACE2), several airway proteases, interferons and interferon-stimulated genes (ISGs), cytokines, transcription factors, pathogen recognition receptors, mucins, and surfactants. We report here that sex significantly affects hNEC transcriptional responses to SARS-CoV-2 infection and to WSP exposure. We found that males displayed a more robust upregulation of immune and antiviral genes in response to infection compared to females, but WSP exposure prior to infection significantly downregulated these genes in females with few effects in males. Together, data presented here provide a link between exposure to WSP and modification of SARS-CoV-2 induced antiviral host defense responses in the nasal mucosa.

## Methods

### Primary Nasal Epithelial Cell Donors

Collection of primary hNECs from adults was performed as previously described (22). Superficial nasal epithelial scrape biopsies were obtained from N=13 (7F, 6M) healthy, non-smoking adults with a Rhino-Pro curette (Arlington Scientific, Inc. 96-0900) per protocols approved by the University of North Carolina School of Medicine Institutional Review Board for Biomedical Research (protocol numbers 05-2528, 09-0716, 11-1363). Written informed consent was obtained from all study participants. Demographic information about the donors used for each exposure including age, BMI, and race is provided in Table 1. Nasal biopsies were stored in RPMI-1640 medium (Gibco 11875-093) on ice until further processing.

**Table 1:**
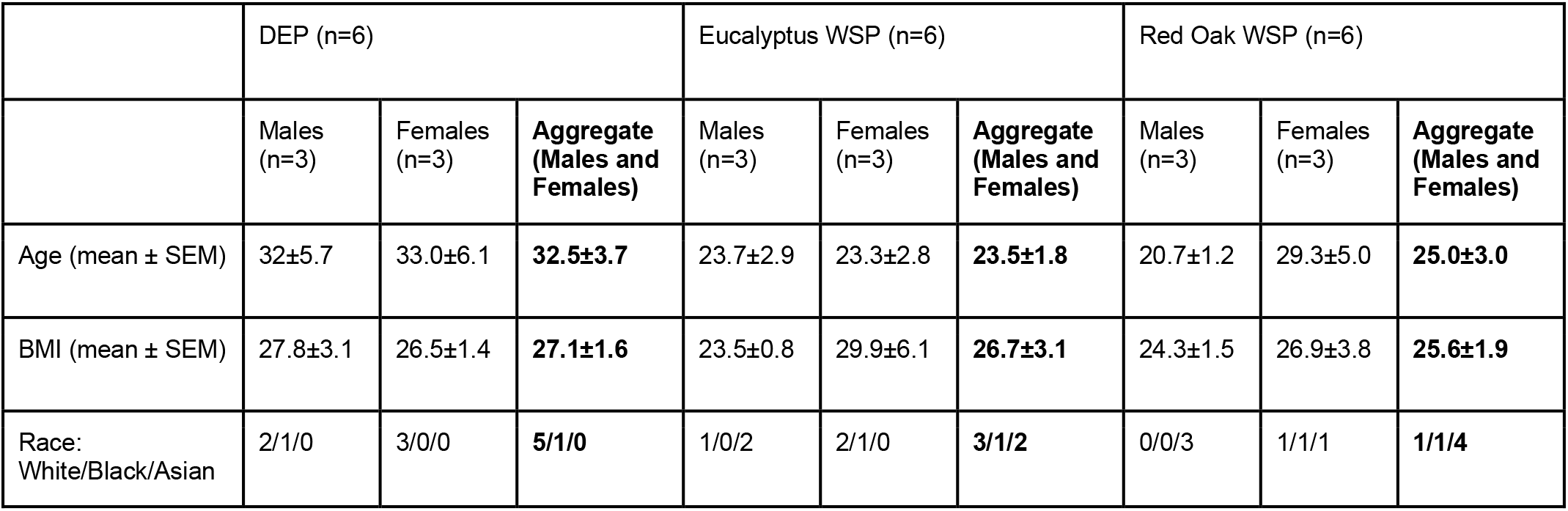
Demographic information about hNEC donors

|  | DEP (n=6) |  |  | Eucalyptus WSP (n=6) |  |  | Red Oak WSP (n=6) |  |  |
| --- | --- | --- | --- | --- | --- | --- | --- | --- | --- |
|  | Males<br>(n=3) | Females<br>(n=3) | Aggregate<br>(Males and<br>Females) | Males<br>(n=3) | Females<br>(n=3) | Aggregate<br>(Males and<br>Females) | Males<br>(n=3) | Females<br>(n=3) | Aggregate<br>(Males and<br>Females) |
| Age (mean ± SEM) | 32±5.7 | 33.0±6.1 | 32.5±3.7 | 23.7±2.9 | 23.3±2.8 | 23.5±1.8 | 20.7±1.2 | 29.3±5.0 | 25.0±3.0 |
| BMI (mean ± SEM) | 27.8±3.1 | 26.5±1.4 | 27.1±1.6 | 23.5±0.8 | 29.9±6.1 | 26.7±3.1 | 24.3±1.5 | 26.9±3.8 | 25.6±1.9 |
| Race:<br>White/Black/Asian | 2/1/0 | 3/0/0 | 5/1/0 | 1/0/2 | 2/1/0 | 3/1/2 | 0/0/3 | 1/1/1 | 1/1/4 |

### Expansion and Culture of hNECs

Culture of hNECs was performed as previously described (22, 27). Cells from nasal biopsy were expanded at passage 0 on a 12-well, PureCol-coated (Advanced Biomatrix 5005-100ML) cell culture plate (Costar 3512) in PneumaCult -Ex Plus Medium (Stemcell Technologies 05041, 05042) supplemented with hydrocortisone (Stemcell Technologies 07925), antibiotic antimycotic solution (Sigma A5955), and gentamicin reagent solution (Gibco 15750-060). Cells were passaged and further expanded in 25 cm^2^ tissue culture flasks (Corning 430639) until passage 2. HNECs were then seeded on 12 mm transwell inserts with 0.4 μm pores (Costar 3460) coated with human placental collagen (Sigma C7521-10MG) at a density of 203,000-333,000 cells per well and maintained in PneumaCult -Ex Plus Medium. Once confluency was reached on the transwells, the cultures were taken to air-liquid-interface (ALI) and the apical medium was permanently removed, while the basolateral medium was switched for PneumaCult ALI Medium (Stemcell Technologies 05002, 05003, 05006), supplemented with 1% pen strep (Gibco 15140-122), hydrocortisone (Stemcell Technologies 07925), and heparin (Stemcell Technologies 07980). After this point, three times per week the basolateral medium was changed and the apical surfaces of the cultures were washed with 37°C HBSS + CaCl_2_, + MgCl_2_ (HBSS++) (Gibco 14025-092). Mucociliary differentiation of the cultures was achieved after 4-6 weeks of ALI conditions. At the time of exposure, cultures were at ALI for 5.29-9.14 weeks.

### Diesel Exhaust Particulate (DEP) Suspension Preparation

Whole diesel exhaust particulate material from an automobile engine was collected as described by Sagai, et al. (39). Twenty-five mg of the DEP was diluted in 5 ml of warmed (37°C) phenol red-free MEM basal medium (Gibco 51200-038). The suspension was sonicated with a Fisher Sonic Dismembrator Model 500 with a microprobe tip for two 1-minute cycles. During each cycle the probe was moved up and down in the suspension, and sonication alternated between 30% output for 0.5 s and 0% output for 0.5 s. After each cycle, the suspension was mixed by inversion. An additional 20 ml of warmed (37°C) medium was then added to the suspension to achieve a final concentration of 1 mg/ml. Aliquots of the suspension were snap frozen in liquid nitrogen and stored at −80°C for future use.

### Woodsmoke Particulate (WSP) Suspension Preparation

Woodsmoke generated from eucalyptus and red oak were each collected as previously described by Kim, et al. (7). Briefly, eucalyptus or red oak were burned in a quartz tube furnace at 640°C and smoke was collected in a series of cryogenic traps. The resulting woodsmoke particulate condensates were then collected in acetone and concentrated with a rotary evaporator. Finally, the particulates were dried and the solid PM was resuspended in Dulbecco’s PBS (Gibco 14200-075) at 2 mg/ml and frozen at −20°C. Prior to exposure, aliquots were sonicated in a water bath sonicator (Sinosonic Industrial Co. Ltd., Taiwan, Model B200) at 40 KHz for 4.75 minutes.

### Particle Size Measurements of Particulate Suspensions

Particle size distributions of the three particulate suspensions were determined by diluting an aliquot of each to 50 μg/ml in ddH_2_O. The diluted suspensions were run through a BD FACSVerse 2013 Flow Cytometer for size measurement and compared to size calibration standards (Thermo Fisher F13838) of 1.0, 2.0, 4.0, 6.0, 10.0, and 15.0 μm in diameter. Graphs of particle size distribution overlaid with the standard sizes are shown in [Supplemental Fig. S1](https://figshare.com/s/6ba4f3191158622d0d8e).

### Exposure of hNECs to DEP or WSP

A pictorial depiction of the exposure and infection scheme is provided in Fig. 2. The apical surface of each culture was washed with 100 μl of warmed (37°C) HBSS++ and carefully aspirated. Basolateral medium was removed and replaced with 1.0 ml of 37°C PneumaCult ALI Medium. Warm ALI Medium was used as the control exposure and as the vehicle for particulate exposures. HNECs from three male and three female donors were used for each type of exposure (DEP, eucalyptus WSP, and red oak WSP). Particulate stock aliquots were diluted in ALI medium and applied to the apical surface of the experimental wells at a concentration of 22 μg/cm^2^ in 150 μl, a dose we have studied previously (24). Control wells received 150 μl ALI medium apically. Cultures were then returned to the incubator (37°C, 5% CO_2_) for 2 h.

**Fig. 2.**
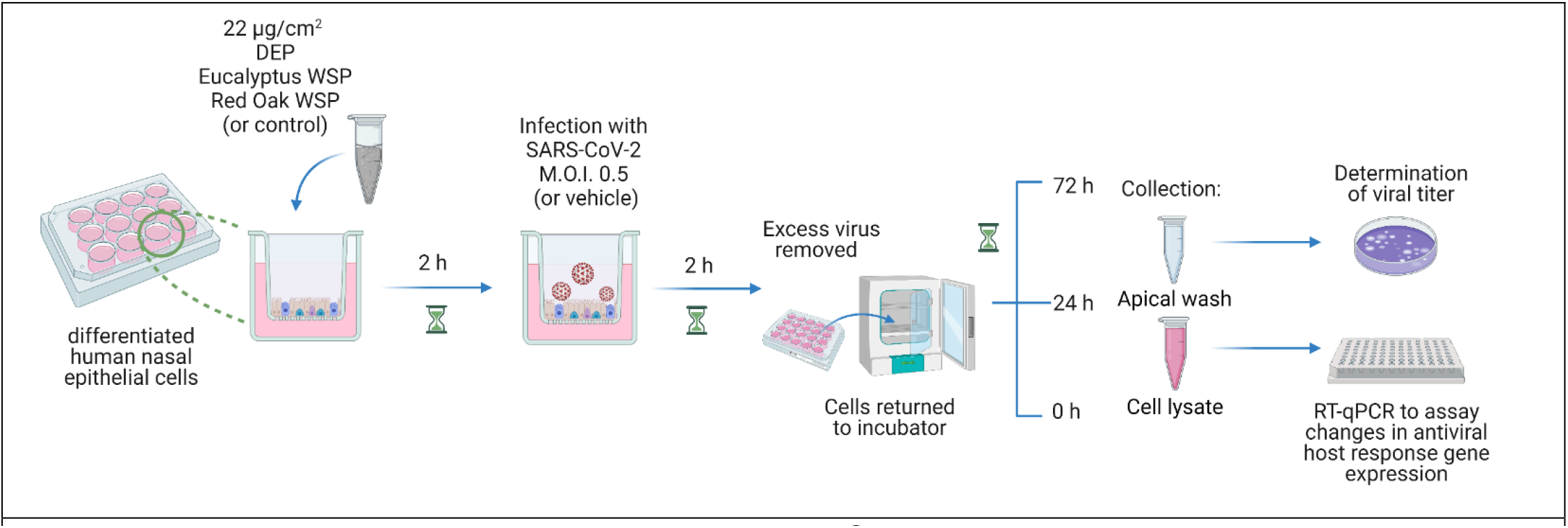
Experimental design scheme. Differentiated hNECs from males and females grown at ALI were exposed to 22 μg/cm^2^ DEP, eucalyptus WSP, or red oak WSP (or control) for 2 h. At the end of the exposure period, cells were infected with SARS-CoV-2 at an M.O.I. of 0.5 (or mock infected with vehicle) for 2 h. Excess virus/vehicle was then removed and the apical surface was washed. A second apical wash and cell lysis were performed immediately or 24 or 72 h later. Apical washes were used to determine viral titers and RNA was purified from cell lysates and used for RT-qPCR to assess altered gene expression in a panel of 48 genes. Figure created with BioRender.com.

### Infection with SARS-CoV-2 or Mock

At the end of the 2-h exposure, half the wells exposed to particulate and half the control wells were apically infected with SARS-CoV-2 derived from clinical isolate WA1 (20) in high glucose DMEM (Gibco 11995-065) with 5% heat-inactivated fetal bovine serum, 1% L-glutamine diluted in ALI medium at a M.O.I. (multiplicity of infection) of 0.5 in 100 μl. The other half of the cultures were mock infected with 100 μl of high glucose DMEM with 5% heat-inactivated fetal bovine serum, 5% L-glutamine diluted in ALI medium. Cultures were then returned to the incubator (37°C, 5% CO_2_) for 2 h.

### Sample Collection

After the 2-h infection, cells were checked under the microscope for signs of cell death. The apical liquid was carefully removed from every well. Cultures were then washed with 200 μl 37°C HBSS++ and returned to the incubator until collection. At the time of collection (0, 24, or 72 h p.i.), 100 μl 37°C HBSS++ was added to the apical surface of each culture, and cells were returned to the incubator for 15 m. Apical washes were then carefully collected and analyzed for viral titer. Cells were lysed using 350 μl cold TRIzol reagent (Life Technologies 15596018) for subsequent gene expression analysis.

### Determination of Viral Titer

Fifty microliters of the apical wash were mixed with 450 μl of medium (DMEM + 5% FBS + 1% L-glutamine) followed by ten-fold serial dilutions resulting in a dilution series of 10^-1^ to 10^-6^. Two hundred μl of each dilution was added to plated Vero E6 cells (C1008, ATCC) and incubated at 37°C. Plates were rocked every 15 min to ensure even distribution of the virus over the surface of the well. After 1 h, 2 ml of overlay (50:50 mixture of 2.5% carboxymethylcellulose and 2X alpha MEM containing 6% FBS + 2% penicillin/streptomycin + 2% L-glutamine + 2% HEPES) was added to each well. Plates were incubated at 37°C, 5% CO_2_ for 4 days, then fixed with 2 ml of 4% paraformaldehyde left on overnight. Following removal of the fixative, wells were rinsed with water to remove residual overlay and then stained with 0.25% crystal violet. Visible plaques were counted and averaged between two technical replicate wells. Viral titers were calculated as plaque forming units (pfu) per ml. The limit of detection for the assay was determined to be 12.5 pfu / wash, and samples that yielded no plaques were assigned a value of 6.25, half of the limit of detection.

### RNA extraction from whole cell lysates in TRIzol

Whole cell lysates in TRIzol reagent were thawed on ice. An additional 650 μl cold TRIzol was added to each sample to facilitate RNA collection. Two hundred μl chloroform was added to each tube and tubes were shaken vigorously and incubated at room temperature for 3 minutes. Samples were then centrifuged for 15 minutes at 12,000 x g at 4°C. The aqueous phase containing RNA was then carefully removed from each sample and transferred to new microcentrifuge tubes. One volume of 100% ethanol was added per volume of aqueous phase removed and samples were vortexed. Samples were further processed with the Zymo RNA Clean and Concentrator Kit (Zymo R1016) according to the manufacturer’s instructions. Eluted RNA was stored at −80°C until use.

### Generation of cDNA and Quantification of Gene Expression of 48 genes by qPCR

RNA concentration and purity were measured using a CLARIOstar plate reader and an LVis Plate (BMG LABTECH). For each sample, 800 ng of RNA was used to generate cDNA in a reaction volume of 25 μl. The final concentrations of reagents in each reaction were as follows: 0.50 mM dNTPs (Promega U151B), 1.00 U/μl RNasin Ribonuclease Inhibitor (Promega N211A), 10.0 U/μl M-MLV Reverse Transcriptase (Invitrogen 28025-013), 0.10 μg/μl Random Primers (Invitrogen 58875), 50.0 mM KCl, 0.25 mM MgCl_2_, 20.0 mM Tris-HCl, 0.01 mg/ml BSA. PCR was performed in 96-well plates (Thermo AB-0600, AB-0851) for one cycle (25.0°C for 10 minutes, 37.0°C for 50 minutes, 70.0°C for 15 minutes, followed by 4.0°C infinite hold). Samples were submitted to the UNC School of Medicine Center for Gastrointestinal Biology and Disease Advanced Analytics Core for high-throughput qPCR gene expression analysis. Gene expression of a panel of 48 genes (including 2 reference genes) was assayed in a Fluidigm BioMark HD system using TaqMan primers and probes. The list of primer/probe catalog numbers for all genes assayed is provided in Table 2. Duplicate Ct values were measured for each sample/gene combination and averaged for further analysis. Gene expression was calculated using the ΔΔCt method with normalization to the geometric mean of expression of the two reference genes (*ACTB* and *GAPDH*). Two samples (out of 216) showing poor amplification across the panel (i.e. comparable to the no-template controls) were excluded from the data set and not further analyzed.

**Table 2:** Genes assayed, grouped by functional categories. Assay identifiers are listed for TaqMan primer/probe sets purchased from Thermo Fisher (or IDT where indicated).

| Gene Name: | Category: | TaqMan Probe Assay ID: |
| --- | --- | --- |
| ACE2 | Viral Entry Factor (VEF) | Hs01085331_m1 |
| CTSB | Airway Proteases | Hs00157194_m1 |
| CTSL |  | Hs00964651_m1 |
| FURIN |  | Hs06637404_sH |
| MMP7 |  | Hs01042796_m1 |
| ST14 |  | Hs01058386_m1 |
| TMPRSS11D |  | Hs00975370_m1 |
| TMPRSS2 |  | Hs05024838_m1 |
| IFIT1 | Antiviral Defense | Hs03027069_s1 |
| IFITM3 |  | Hs03057129_s1 |
| IFNA1 |  | Hs04189288_g1 |
| IFNA2 |  | Hs00265051_s1 |
| IFNB1 |  | Hs00265051_s2 |
| IFNG |  | Hs00265051_s3 |
| IFNL1 |  | Hs00265051_s4 |
| IFNL2 |  | Hs00265051_s5 |
| LTF |  | Hs00265051_s6 |
| MX1 |  | Hs00265051_s7 |
| SOCS3 |  | Hs00265051_s8 |
| CCL2 | Cell Signaling/Immune Cell Recruitment | Hs00265051_s9 |
| CCL3 |  | Hs00265051_s10 |
| CCL5 |  | Hs00265051_s11 |
| CSF2 |  | Hs00265051_s12 |
| CXCL10 |  | Hs00265051_s13 |
| CXCL11 |  | Hs00265051_s14 |
| CXCL8 |  | Hs00265051_s15 |
| CXCL9 |  | Hs00265051_s16 |
| IL15 |  | Hs00265051_s17 |
| IL1B |  | Hs00265051_s18 |
| IL6 |  | Hs00265051_s19 |
| TNF |  | Hs00265051_s20 |
| MUC5AC | Mucins | Hs00265051_s21 |
| MUC5B |  | Hs00265051_s22 |
| SFTPA1 | Surfactant (Surf.) | Hs00265051_s23 |
| SFTPD |  | Hs00265051_s24 |
| IRF1 | Transcription Factors | Hs00265051_s25 |
| IRF3 |  | Hs00265051_s26 |
| IRF7 |  | Hs00265051_s27 |
| NFKB1 |  | Hs00265051_s28 |
| STAT1 |  | Hs00265051_s29 |
| STAT2 |  | Hs00265051_s30 |
| STAT3 |  | Hs00265051_s31 |
| DDX58 | Viral Recognition | Hs00265051_s32 |
| TLR3 |  | Hs00265051_s33 |
| TLR7 |  | Hs00265051_s34 |
| TLR9 |  | Hs00265051_s35 |
| nCoVn1 | Viral Genes | IDT Cat # 10006713 |
| nCoVn2 |  | IDT Cat # 10006713 |
| GAPDH | Reference Genes | Hs00265051_s36 |
| ACTB |  | Hs00265051_s37 |
| RP |  | IDT Cat # 10006713 |

### Quantification of SARS-CoV-2 N1 and N2 gene expression by qPCR

Expression of viral SARS-CoV-2 N1 and N2 genes was also quantified and normalized to human RNase P gene expression using the Integrated DNA Technologies 2019-nCoV RUO Kit (IDT 10006713). For a single reaction, 6.5 μl nuclease-free water, 1.5 μl of one primer/probe mix, and 10 μl of TaqMan Universal Master Mix II, with UNG (Thermo Fisher 4440038) were mixed and added to every well of a Sapphire 96-well PCR Microplate (Greiner Bio-one 652260). cDNA was then added to each well (2 μl) for a total volume of 20 μl per reaction. The plate was sealed with a plate film (Thermo Fisher 4311971) and centrifuged for 5 minutes at 500 x g at room temperature. RT-qPCR was performed on a QuantStudio 3 using the following reaction conditions: hold 50.0°C for 2 minutes then hold 95.0°C for 10 minutes, cycle through 95.0°C for 15 s and 60.0°C for 1 minute for 40 cycles. Transitions between temperatures occurred at 1.6°C/s. The two samples excluded from the Fluidigm PCR data were also excluded here. Results were collected as Ct and analyzed with the ΔΔCt method, normalized to expression of human RNase P.

### Statistical Analysis

Analysis was carried out using SAS PROC MIXED as a full factorial design, with sex (M or F), particulate treatment (control, DEP, eucalyptus WSP, or red oak WSP), virus or no virus, and duration (0, 24, or 72 h), as well as all their interactions. Donor was fit as a random effect. Preplanned hypothesis tests for differences between marginal means were carried out as t-tests with the LSMESTIMATE command. Sex-specific means were calculated for each combination of particulate treatment, virus, and duration and differences were tested using a t-test with the LSMESTIMATE command. Correction for multiple comparisons was performed across all statistical tests for the entire experiment using the ‘qvalue’ R package (v. 2.22.0), with a false discovery rate q-value threshold of 0.05, assuming pi0 = 1 (equivalent to Benjamini-Hochberg correction). The resultant p-value for statistical significance was p ≤ 0.00369. Viral titer data was analyzed using GraphPad Prism v. 8.4.0. Unpaired t-tests (with Welch’s correction when appropriate) were used to evaluate differences in log_10_-transformed data.

## Results

### Particulate exposure does not affect viral load in hNECs

Previously, we found that exposing hNECs and other airway epithelial cells to DEP prior to infection with influenza A enhanced viral replication and susceptibility to viral infection (24). In the present study, we thus sought to determine whether DEP and WSP would have similar effects in a SARS-CoV-2 infection model. HNECs were exposed to control (ALI medium) or 22 μg/cm^2^ DEP, eucalyptus WSP, or red oak WSP for 2 h, followed by infection with SARS-CoV-2. Viral loads in apical washes for the hNECs exposed to particulates and their respective controls at 0, 24, and 72 h p.i. are shown in Fig. 3 A-C. The amount of infectious virus recovered from apical washes increased with duration of infection (Fig. 3 D), suggesting increased viral replication and apical secretion over time, consistent with our previous study (20). Exposure to particulates, regardless of type, had no effect on viral loads in apical washes (Fig. 3 A-C). On average, viral load recovered from males was higher than viral load recovered from hNECs from female donors, though the difference in viral loads between the sexes did not reach statistical significance (Fig. 3 D).

**Fig. 3:**
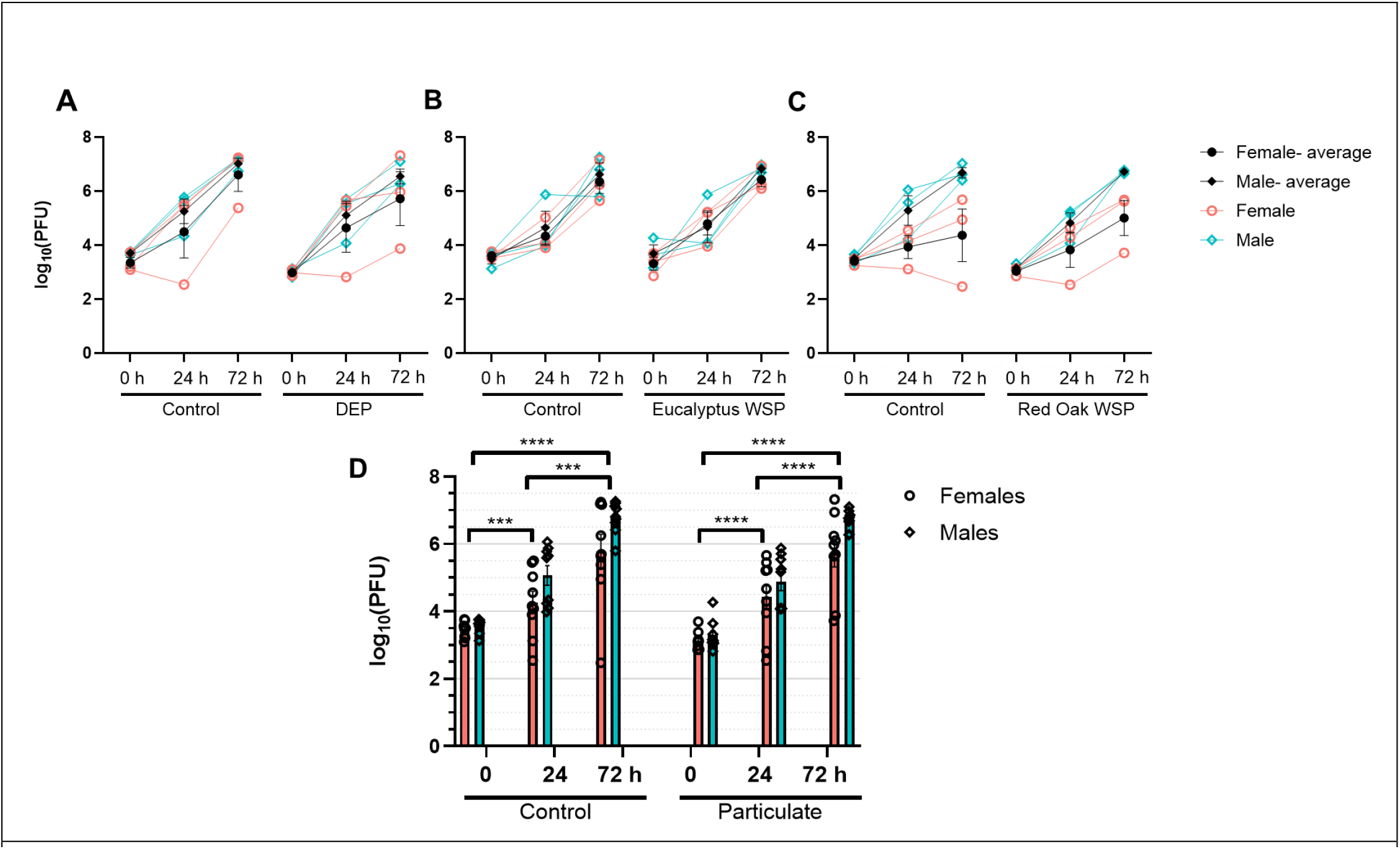
SARS-CoV-2 viral titers in hNEC cultures at 0, 24, and 72 h p.i. hNECs from male and female donors were exposed to particulates (DEP or WSP from flaming eucalyptus or red oak, at 22 μg/cm^2^) or control for 2 h, then infected with SARS-CoV-2 at an MOI of 0.5. At 0, 24, or 72 h post infection, the apical washes were collected and used for approximating viral titer. Titers from individual particulate exposures with respective controls for DEP, eucalyptus WSP, and red oak WSP are shown in **A-C** respectively. Black symbols indicate sex-specific means with standard error bars (N=3 biological replicates each for males and females). **D** Aggregated viral titers recovered from hNECs exposed to vehicle or a particulate. Standard error is shown (N=9 biological replicates for each bar). Unpaired t-tests with Welch’s correction were used to determine (sex aggregated) differences in viral titer between time points, with *** p = 0.0001, **** p < 0.0001.

Expression levels for a panel of 48 genes were determined in hNECs infected with SARS-CoV-2. The relationship between the expression level of each gene (relative to reference genes) and the viral titer recovered from respective samples is shown in Fig. 4. As expected, expression levels of SARS-CoV-2 N1 and N2 genes are highly correlated with viral load recovered (Pearson’s *r* = 0.91 for both). This indicates that apical release of infectious viral particles is highly correlated with viral mRNA levels. The following genes are also correlated with viral titer, with a statistically significant Pearson’s *r* > 0.70: *ACE2, IFIT1, IFITM3, IFNB1, IFNL1, IFNL2, MX1, CCL5, CXCL10, IRF7, STAT1, DDX58*, and *TLR9*. Interestingly, *TMPRSS2* and *IL1B* both appear to be negatively correlated with viral titer.

**Fig. 4:**
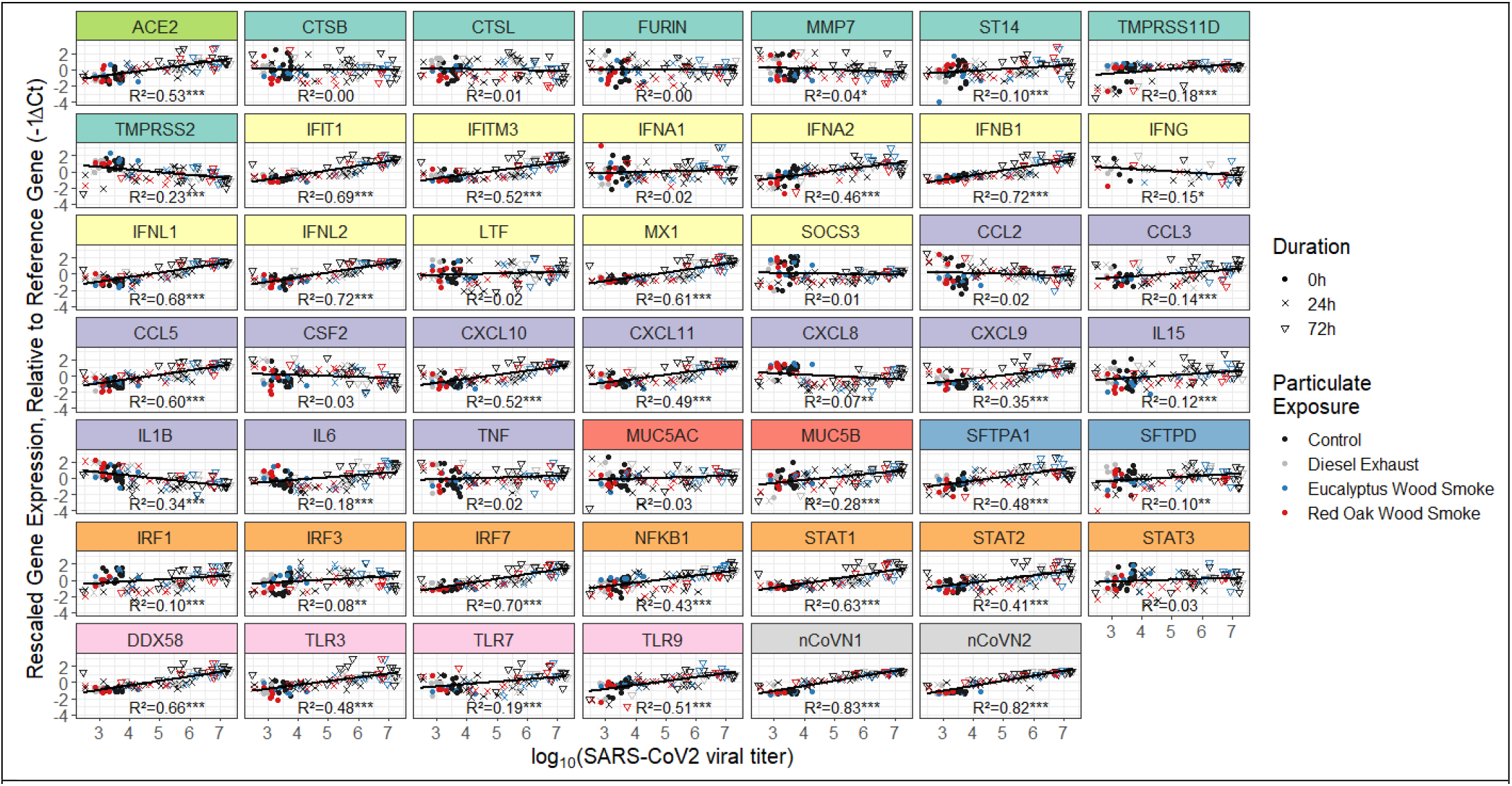
Relationship between gene expression relative to reference genes (-ΔCt) and log_10_(viral titer) in infected cells. Colors behind gene names correspond to functional categories presented in Table 2. Statistical significance is indicated next to the coefficient of determination (R^2^): * p ≤ 0.05, ** p ≤ 0.01, *** p ≤ 0.001.

### SARS-CoV-2 infection greatly affects expression of antiviral host response genes in hNECs

In order to assess how particulate exposure affects expression of antiviral host defense genes in the presence of an infection, we first needed to measure the independent effects of SARS-CoV-2 infection on gene expression. Thus, hNECs from male and female donors which were not exposed to any particulates were infected with SARS-CoV-2 (or mock infected with vehicle). At 0, 24, or 72 h p.i., cells were lysed, and RNA was extracted and purified for RT-qPCR. Gene expression in infected cultures was compared to that of corresponding uninfected cultures after normalization to reference gene expression. Virus-induced changes in gene expression in hNECs at 0, 24, and 72 h p.i. are shown in Fig. 5 and fold-inductions and p-values are tabulated in Table 3. By 24 h p.i., the Type III IFNs (*IFNL1* and *IFNL2*) were upregulated in hNECs from male and female donors, with statistically significant upregulation of both genes in males. Expression of *IFNL1* and *IFNL2* were even more highly upregulated at 72 h p.i. and reached statistical significance in both sexes. Additionally, by 72 h p.i., many other genes related to antiviral defense, cell signaling, and immune cell recruitment were significantly upregulated relative to the uninfected cells. Transcription factors, especially *IRF7* and *STAT1* were upregulated in hNECs from male and female donors at 72 h p.i., and *DDX58*, which encodes RIG-I, a cytoplasmic viral nucleic acid receptor, was also upregulated in both sexes. *ACE2* expression was significantly upregulated at 72 h p.i. in hNECs from both sexes. In most instances, gene expression in hNECs from males was more highly induced by infection than in hNECs from females, suggesting an overall more robust epithelial response to SARS-CoV-2 in hNECs from male donors. For each gene that was differentially expressed in infected cells from both sexes, the ratio of expression in males:females was calculated. Indeed, on average the level of virus-induced gene expression in hNECs from males was 2.08 times (95% CI: ± 0.57) that of hNECs from females. Additionally, we assessed whether baseline differences in gene expression existed between the sexes in uninfected cells. There were no statistically significant differences in baseline gene expression between hNECs from males and females at 24 and 72 h post mock infection (data not shown).

**Fig. 5:**
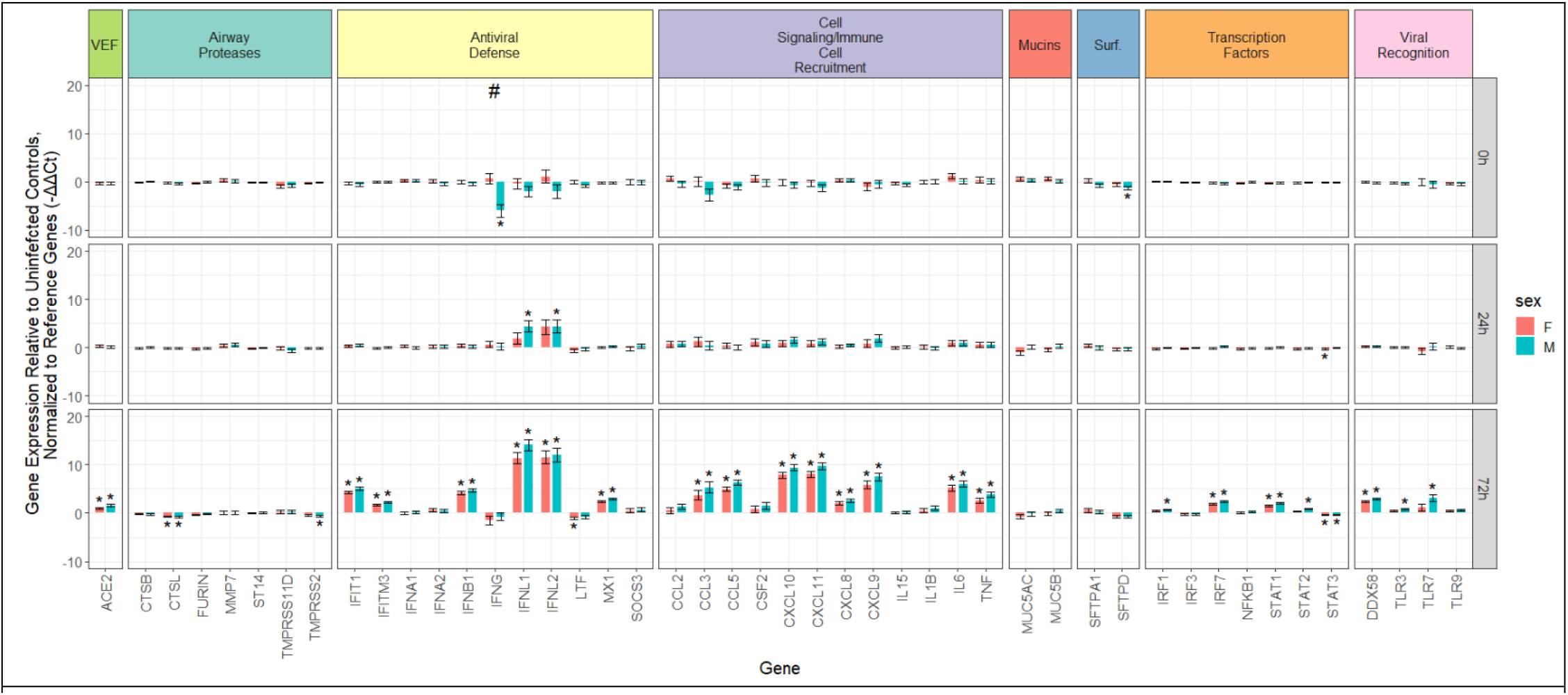
Gene expression in infected hNECs from males and females relative to uninfected controls (-ΔΔCt) at 0, 24, and 72 h p.i. Gene categories are color-coded at the top, with ‘VEF’ an abbreviation for ‘Viral Entry Factor’ and ‘Surf.’ an abbreviation for ‘Surfactant’. Graphed as average (N=9 biological replicates for each bar) with standard error. Statistically significant (q ≤ 0.05) changes in gene expression are represented by *. A statistically significant difference in gene expression between males and females is indicated by #.

**Table 3:**
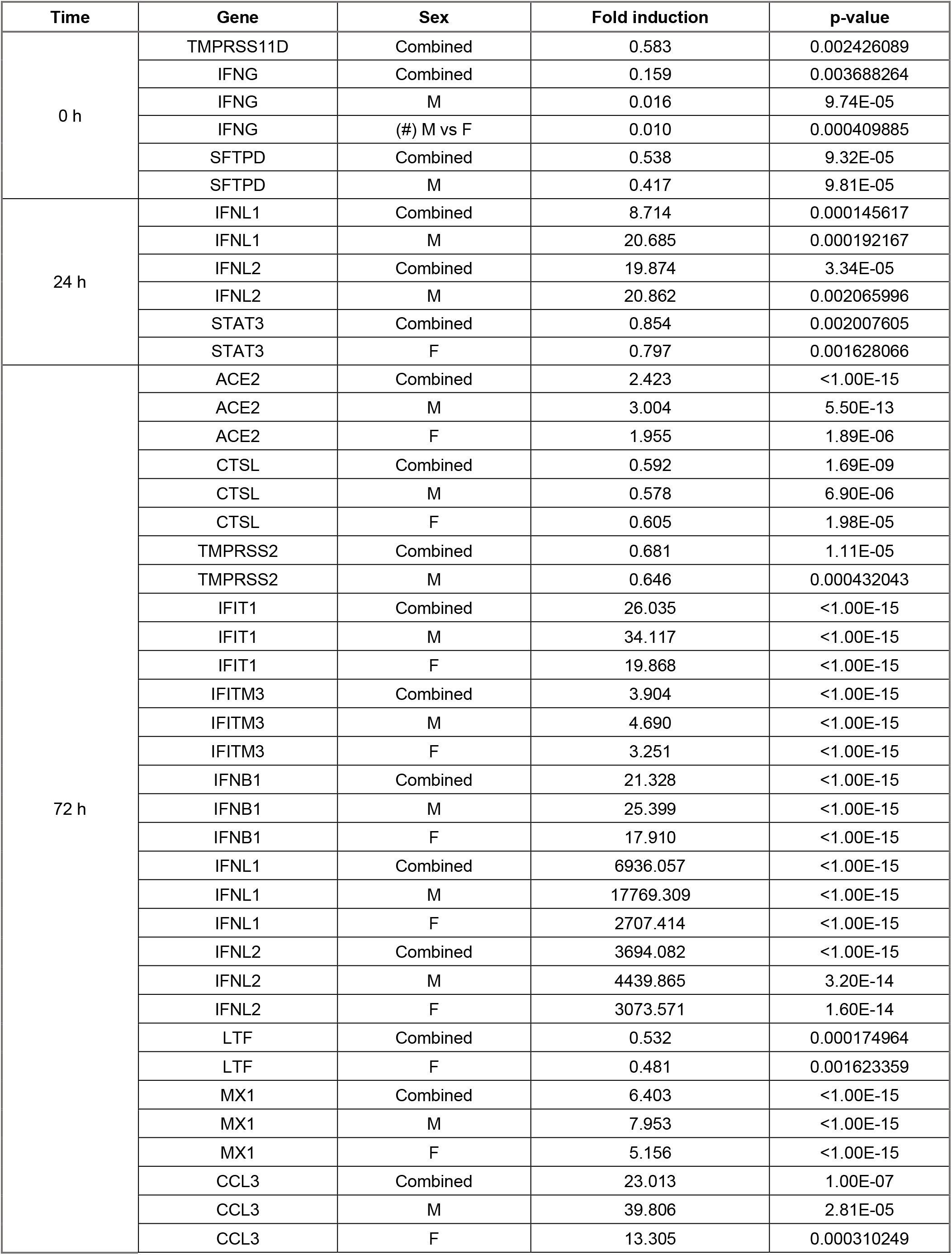
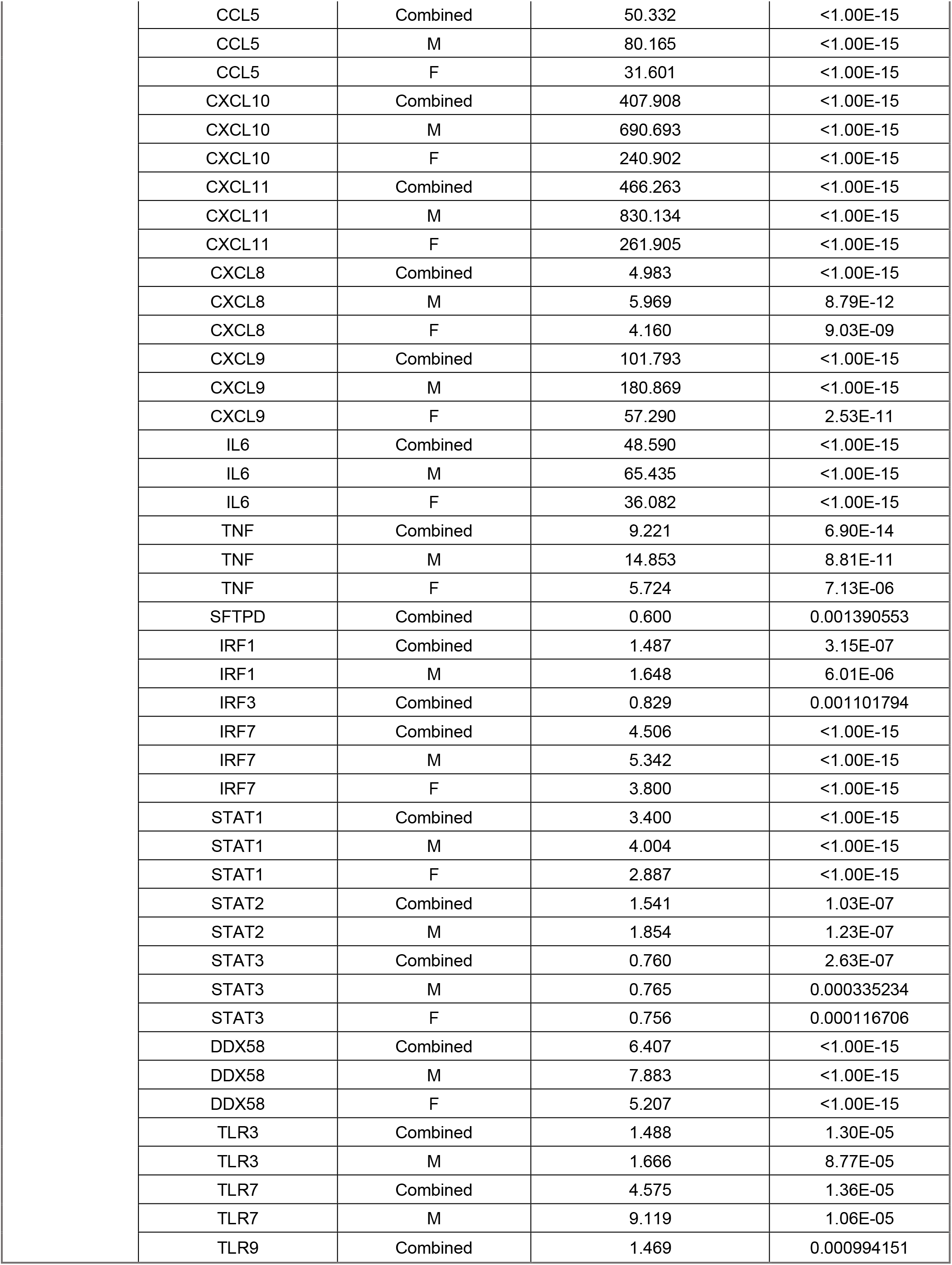
Statistically significant virus-induced changes in gene expression in hNECs from males and females at 0, 24, and 72 h p.i. with SARS-CoV-2. N=9 biological replicates for individual sex effects (M or F) and N=18 for Combined effects.

| Time | Gene | Sex | Fold induction | p-value |
| --- | --- | --- | --- | --- |
| 0 h | TMPRSS11D | Combined | 0.583 | 0.002426089 |
|  | IFNG | Combined | 0.159 | 0.003688264 |
|  | IFNG | M | 0.016 | 9.74E-05 |
|  | IFNG | (#) M vs F | 0.010 | 0.000409885 |
|  | SFTPD | Combined | 0.538 | 9.32E-05 |
|  | SFTPD | M | 0.417 | 9.81E-05 |
| 24 h | IFNL1 | Combined | 8.714 | 0.000145617 |
|  | IFNL1 | M | 20.685 | 0.000192167 |
|  | IFNL2 | Combined | 19.874 | 3.34E-05 |
|  | IFNL2 | M | 20.862 | 0.002065996 |
|  | STAT3 | Combined | 0.854 | 0.002007605 |
|  | STAT3 | F | 0.797 | 0.001628066 |
| 72 h | ACE2 | Combined | 2.423 | <1.00E-15 |
|  | ACE2 | M | 3.004 | 5.50E-13 |
|  | ACE2 | F | 1.955 | 1.89E-06 |
|  | CTSL | Combined | 0.592 | 1.69E-09 |
|  | CTSL | M | 0.578 | 6.90E-06 |
|  | CTSL | F | 0.605 | 1.98E-05 |
|  | TMPRSS2 | Combined | 0.681 | 1.11E-05 |
|  | TMPRSS2 | M | 0.646 | 0.000432043 |
|  | IFIT1 | Combined | 26.035 | <1.00E-15 |
|  | IFIT1 | M | 34.117 | <1.00E-15 |
|  | IFIT1 | F | 19.868 | <1.00E-15 |
|  | IFITM3 | Combined | 3.904 | <1.00E-15 |
|  | IFITM3 | M | 4.690 | <1.00E-15 |
|  | IFITM3 | F | 3.251 | <1.00E-15 |
|  | IFNB1 | Combined | 21.328 | <1.00E-15 |
|  | IFNB1 | M | 25.399 | <1.00E-15 |
|  | IFNB1 | F | 17.910 | <1.00E-15 |
|  | IFNL1 | Combined | 6936.057 | <1.00E-15 |
|  | IFNL1 | M | 17769.309 | <1.00E-15 |
|  | IFNL1 | F | 2707.414 | <1.00E-15 |
|  | IFNL2 | Combined | 3694.082 | <1.00E-15 |
|  | IFNL2 | M | 4439.865 | 3.20E-14 |
|  | IFNL2 | F | 3073.571 | 1.60E-14 |
|  | LTF | Combined | 0.532 | 0.000174964 |
|  | LTF | F | 0.481 | 0.001623359 |
|  | MX1 | Combined | 6.403 | <1.00E-15 |
|  | MX1 | M | 7.953 | <1.00E-15 |
|  | MX1 | F | 5.156 | <1.00E-15 |
|  | CCL3 | Combined | 23.013 | 1.00E-07 |
|  | CCL3 | M | 39.806 | 2.81E-05 |
|  | CCL3 | F | 13.305 | 0.000310249 |

|  |  |  |  |
| --- | --- | --- | --- |
| CCL5 | Combined | 50.332 | <1.00E-15 |
| CCL5 | M | 80.165 | <1.00E-15 |
| CCL5 | F | 31.601 | <1.00E-15 |
| CXCL10 | Combined | 407.908 | <1.00E-15 |
| CXCL10 | M | 690.693 | <1.00E-15 |
| CXCL10 | F | 240.902 | <1.00E-15 |
| CXCL11 | Combined | 466.263 | <1.00E-15 |
| CXCL11 | M | 830.134 | <1.00E-15 |
| CXCL11 | F | 261.905 | <1.00E-15 |
| CXCL8 | Combined | 4.983 | <1.00E-15 |
| CXCL8 | M | 5.969 | 8.79E-12 |
| CXCL8 | F | 4.160 | 9.03E-09 |
| CXCL9 | Combined | 101.793 | <1.00E-15 |
| CXCL9 | M | 180.869 | <1.00E-15 |
| CXCL9 | F | 57.290 | 2.53E-11 |
| IL6 | Combined | 48.590 | <1.00E-15 |
| IL6 | M | 65.435 | <1.00E-15 |
| IL6 | F | 36.082 | <1.00E-15 |
| TNF | Combined | 9.221 | 6.90E-14 |
| TNF | M | 14.853 | 8.81E-11 |
| TNF | F | 5.724 | 7.13E-06 |
| SFTPD | Combined | 0.600 | 0.001390553 |
| IRF1 | Combined | 1.487 | 3.15E-07 |
| IRF1 | M | 1.648 | 6.01E-06 |
| IRF3 | Combined | 0.829 | 0.001101794 |
| IRF7 | Combined | 4.506 | <1.00E-15 |
| IRF7 | M | 5.342 | <1.00E-15 |
| IRF7 | F | 3.800 | <1.00E-15 |
| STAT1 | Combined | 3.400 | <1.00E-15 |
| STAT1 | M | 4.004 | <1.00E-15 |
| STAT1 | F | 2.887 | <1.00E-15 |
| STAT2 | Combined | 1.541 | 1.03E-07 |
| STAT2 | M | 1.854 | 1.23E-07 |
| STAT3 | Combined | 0.760 | 2.63E-07 |
| STAT3 | M | 0.765 | 0.000335234 |
| STAT3 | F | 0.756 | 0.000116706 |
| DDX58 | Combined | 6.407 | <1.00E-15 |
| DDX58 | M | 7.883 | <1.00E-15 |
| DDX58 | F | 5.207 | <1.00E-15 |
| TLR3 | Combined | 1.488 | 1.30E-05 |
| TLR3 | M | 1.666 | 8.77E-05 |
| TLR7 | Combined | 4.575 | 1.36E-05 |
| TLR7 | M | 9.119 | 1.06E-05 |
| TLR9 | Combined | 1.469 | 0.000994151 |

### Particulate exposure alone has modest effects on expression of antiviral host response genes

Next, we assessed how exposure to particulates alone, without subsequent viral infection, would affect expression of our panel of antiviral host response genes. Again, hNECs from male and female donors were exposed to one of three particulates (DEP, eucalyptus WSP, or red oak WSP) or control for 2 h, followed by a “mock” infection for 2 h. Results are shown graphically in [Supplemental Fig. S2](https://figshare.com/s/6ba4f3191158622d0d8e) and statistically significant results are reported in [Supplemental Table S1](https://figshare.com/s/6ba4f3191158622d0d8e). At 0 h p.i. (mock infection), exposure to both types of WSP increased expression of *IL6* and eucalyptus WSP also upregulated expression of *IL1B*. Further, DEP and red oak WSP significantly decreased expression of *IFNG* at 0 h p.i. (data for eucalyptus WSP not shown due to missing data points). By 24 h p.i. both eucalyptus WSP and red oak WSP further upregulated *IL1B* expression, while *IL6* was no longer upregulated. Overall, by 24 and 72 h p.i., particulate treatment in the absence of infection modestly affected expression of the genes in our panel in hNECs.

### Woodsmoke particulates affect expression of virus-induced genes in hNECs infected with SARS-CoV-2

We hypothesized that exposure to particulates would dampen expression of crucial antiviral host response genes upon subsequent SARS-CoV-2 infection. To test this, hNECs from male and female donors were exposed to control or DEP, eucalyptus WSP, or red oak WSP for 2 h, followed by infection with SARS-CoV-2. Overall, red oak WSP caused more statistically significant changes in virus-induced gene expression than the other particulates (Table 4). DEP had very few effects on gene expression in infected cells. Further, for both types of WSP the number of statistically significant effects increased with duration of infection. More specifically, at 0 h p.i., both types of WSP increased *IL1B* and *IL6* expression compared to unexposed, infected cells, with red oak WSP exposure generating more potent upregulation of *IL6*. By 24 h p.i, all three types of particulates upregulated *IL1B* to similar degrees. At 72 h p.i. WSP exposure, especially from red oak, decreased expression of several genes, including *IFNB1, CCL3, CCL5, CXCL10*, and *CXCL11* (Fig. 6). Red oak WSP also decreased expression of *IFNL1* and *IFNL2*, albeit not statistically significantly. Other genes that are important for the antiviral response were also downregulated by eucalyptus and/or red oak, such as *IFIT1, IFITM3, MX1, IRF7, STAT1, STAT2, DDX58*, and *MMP7*. Thus, exposure to WSP prior to infection with SARS-CoV-2 suppressed IFN-dependent immune gene expression.

**Fig. 6:**
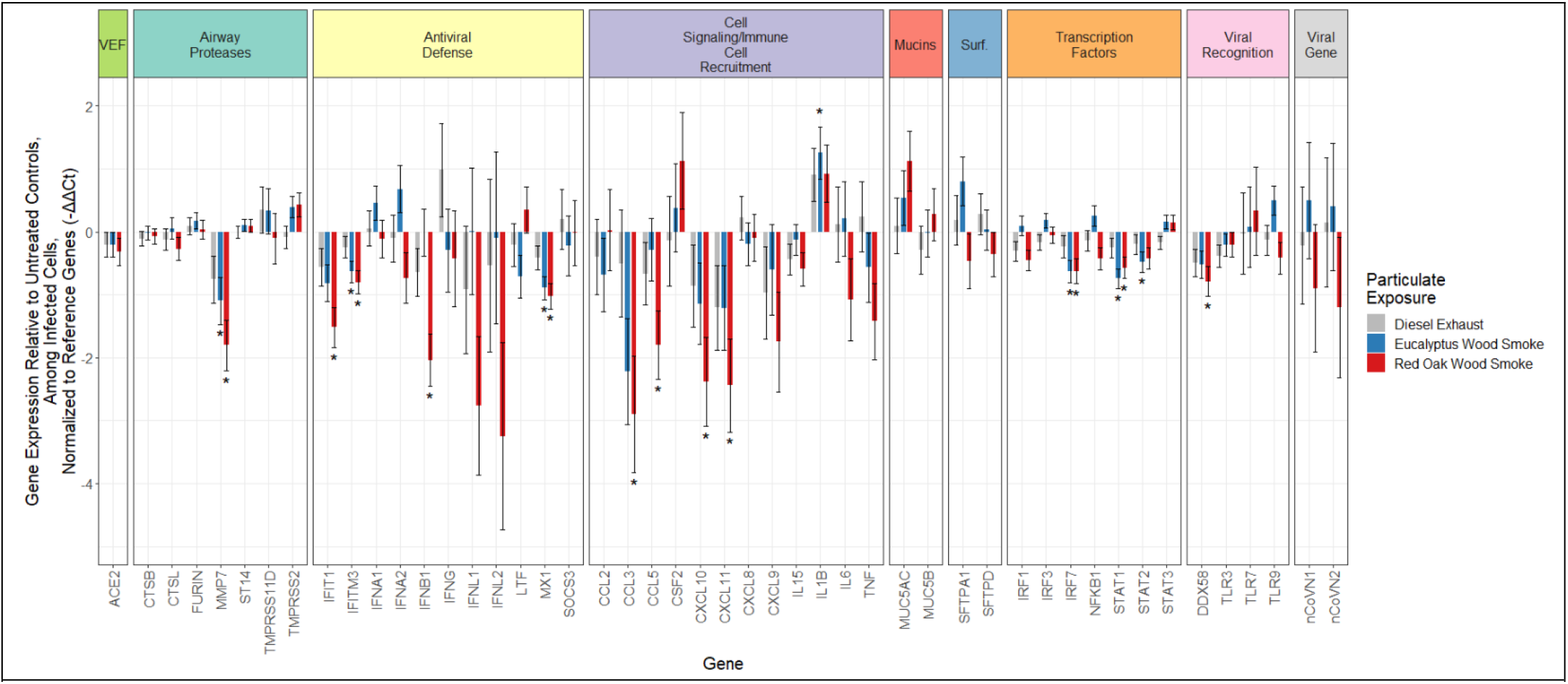
Effects of particulate exposure (DEP and WSP) on virus-induced gene expression in infected hNECs at 72 h p.i. Graphed as means with black bars representing standard error. Males and females are combined for N=6 biological replicates per bar. Statistically significant changes in gene expression are indicated by * (q ≤ 0.05).

**Table 4.** Statistically significant effects of particulate exposure on virus-induced gene expression in hNECs at 0, 24, and 72 h p.i. N=6 (3M, 3F) biological replicates per measurement.

| Time | Gene | Particulate | Sex | Fold induction | p-value |
| --- | --- | --- | --- | --- | --- |
| 0 h | IRF1 | DEP | Combined | 0.638 | 3.60725E-05 |
|  | IL1B | Eucalyptus WSP | Combined | 2.553 | 0.001377971 |
|  | IL6 | Eucalyptus WSP | Combined | 4.024 | 7.84E-04 |
|  | CTSB | Red Oak WSP | Combined | 0.780 | 0.001893867 |
|  | IL1B | Red Oak WSP | Combined | 2.496 | 1.97E-03 |
|  | IL6 | Red Oak WSP | Combined | 8.519 | 5.59E-07 |
|  | STAT2 | Red Oak WSP | Combined | 0.722 | 0.003403266 |
| 24 h | IL1B | DEP | Combined | 3.530 | 2.77506E-05 |
|  | IL1B | Eucalyptus WSP | Combined | 3.691 | 1.13E-05 |
|  | CTSB | Red Oak WSP | Combined | 0.736 | 0.000145452 |
|  | FURIN | Red Oak WSP | Combined | 0.708 | 0.00023035 |
|  | MMP7 | Red Oak WSP | Combined | 0.413 | 0.000723912 |
|  | MX1 | Red Oak WSP | Combined | 0.663 | 1.68E-03 |
|  | IL1B | Red Oak WSP | Combined | 3.299 | 6.47E-05 |
|  | STAT2 | Red Oak WSP | Combined | 0.690 | 8.85E-04 |
| 72 h | MMP7 | Eucalyptus WSP | Combined | 0.467 | 3.38E-03 |
|  | IFITM3 | Eucalyptus WSP | Combined | 0.644 | 2.08E-04 |
|  | MX1 | Eucalyptus WSP | Combined | 0.539 | 3.55E-06 |
|  | IL1B | Eucalyptus WSP | Combined | 2.385 | 3.20E-03 |
|  | IRF7 | Eucalyptus WSP | Combined | 0.647 | 6.52E-04 |
|  | STAT1 | Eucalyptus WSP | Combined | 0.597 | 4.38E-06 |
|  | STAT2 | Eucalyptus WSP | Combined | 0.717 | 2.85E-03 |
|  | MMP7 | Red Oak WSP | Combined | 0.286 | 1.64556E-05 |
|  | IFIT1 | Red Oak WSP | Combined | 0.349 | 5.96E-06 |
|  | IFITM3 | Red Oak WSP | Combined | 0.572 | 2.03E-05 |
|  | IFNB1 | Red Oak WSP | Combined | 0.243 | 2.10E-06 |
|  | MX1 | Red Oak WSP | Combined | 0.493 | 1.46E-06 |
|  | CCL3 | Red Oak WSP | Combined | 0.134 | 2.30E-03 |
|  | CCL5 | Red Oak WSP | Combined | 0.287 | 1.15E-03 |
|  | CXCL10 | Red Oak WSP | Combined | 0.192 | 9.47E-04 |
|  | CXCL11 | Red Oak WSP | Combined | 0.184 | 1.13E-03 |
|  | IRF7 | Red Oak WSP | Combined | 0.647 | 1.90E-03 |
|  | STAT1 | Red Oak WSP | Combined | 0.673 | 1.07E-03 |
|  | DDX58 | Red Oak WSP | Combined | 0.578 | 1.08E-03 |

### Woodsmoke particulate effects on gene expression in infected hNECs are sex-specific

Because the virus-induced effects on gene expression were sex-dependent (Fig. 5), we next assessed whether gene expression changes in cells exposed to particulates prior to virus infection were also sex-dependent. Few sex-specific changes were observed at 0 and 24 h p.i. (Table 5), however at 72 h p.i., WSP, especially red oak, modified virus-induced gene expression in hNECs from female donors (Fig. 7). At this timepoint, WSP from red oak caused statistically significant downregulation of *IFIT1, IFITM3, IFNB1, IRF7, STAT1, DDX58, CXCL10*, and *CXCL11. IFNL1* and *IFNL2* were also downregulated by red oak WSP in hNECs from females but statistical significance was not reached. Additionally, red oak WSP caused a statistically significant decline in *MX1* expression in hNECs from females versus males. Eucalyptus WSP also caused sex-dependent downregulation of several genes in female donors, though the effects were more modest. These results suggest that WSP exposure, especially from red oak, greatly dampens expression of antiviral genes in hNECs from females during SARS-CoV-2 infection, with more modest effects seen in hNECS from males.

**Fig. 7:**
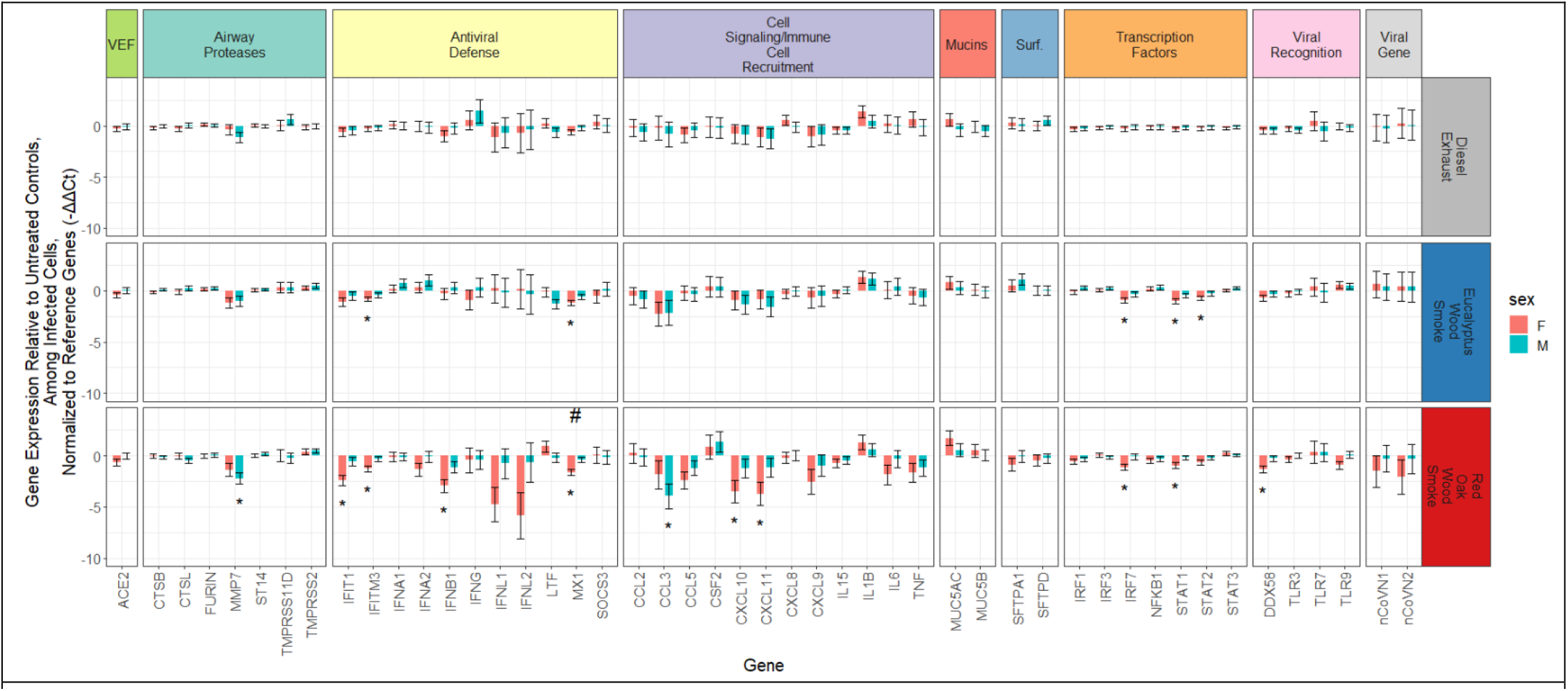
Effects of particulate exposure on virus-induced gene expression in infected hNECs from males or females at 72 h post infection. Graphed as means with black bars representing standard error, N=3 biological replicates per bar. Statistically significant changes in gene expression are represented by * and statistically significant differences in expression between males and females are represented by # (q ≤ 0.05).

**Table 5.** Statistically significant, sex-disaggregated effects of particulate exposure on virus-induced gene expression in infected hNECs at 0, 24, and 72 h p.i. N=3 biological replicates per measurement.

| Time | Gene | Particulate | Sex | Fold induction | p-value |
| --- | --- | --- | --- | --- | --- |
| 0 h | IRF1 | DEP | M | 0.586 | 0.00046212 |
|  | IL6 | Red Oak WSP | M | 8.773 | 0.000240913 |
|  | IL6 | Red Oak WSP | F | 8.271 | 3.87E-04 |
|  | TLR3 | Red Oak WSP | M | 0.580 | 0.002197364 |
| 24 h | IL1B | DEP | M | 4.956 | 1.57E-04 |
|  | IL1B | Eucalyptus WSP | F | 4.695 | 2.17E-04 |
|  | FURIN | Red Oak WSP | F | 0.665 | 0.002092396 |
|  | IL1B | Red Oak WSP | F | 3.398 | 0.003538543 |
| 72 h | IFITM3 | Eucalyptus WSP | F | 0.565 | 5.88E-04 |
|  | MX1 | Eucalyptus WSP | F | 0.434 | 8.25069E-06 |
|  | IRF7 | Eucalyptus WSP | F | 0.524 | 0.000329325 |
|  | STAT1 | Eucalyptus WSP | F | 0.493 | 7.24008E-06 |
|  | STAT2 | Eucalyptus WSP | F | 0.615 | 1.97E-03 |
|  | MMP7 | Red Oak WSP | M | 0.215 | 4.72E-05 |
|  | IFIT1 | Red Oak WSP | F | 0.184 | 1.63E-06 |
|  | IFITM3 | Red Oak WSP | F | 0.413 | 9.25E-06 |
|  | IFNB1 | Red Oak WSP | F | 0.130 | 6.06E-06 |
|  | MX1 | Red Oak WSP | F | 0.322 | 3.90E-07 |
|  | MX1 | Red Oak WSP | (#) M vs F | 2.347 | 2.96E-03 |
|  | CCL3 | Red Oak WSP | M | 0.065 | 1.60E-03 |
|  | CXCL10 | Red Oak WSP | F | 0.089 | 1.35E-03 |
|  | CXCL11 | Red Oak WSP | F | 0.076 | 1.05E-03 |
|  | IRF7 | Red Oak WSP | F | 0.452 | 0.000198042 |
|  | STAT1 | Red Oak WSP | F | 0.516 | 3.37E-04 |
|  | DDX58 | Red Oak WSP | F | 0.403 | 3.60E-04 |

## Discussion

During 2020, air quality reached unhealthy and hazardous levels in the western United States due to wildfires which coincided with the spread of COVID-19. Epidemiological evidence has shown that worsened air quality from PM_2.5_ is associated with increased COVID-19 case rate and case fatality rate around the world. Additionally, toxicological studies have shown that PM treatment affects the host defense response of the airways upon viral infection. In the present study, we hypothesized that exposing hNECs to PM derived from diesel exhaust and woodsmoke, simulating wildfires, would alter the expression of host antiviral response genes upon subsequent infection with SARS-CoV-2. We also hypothesized that these effects would be sex-dependent. We found that exposure to WSP derived from red oak significantly affected gene expression in the context of SARS-CoV-2 infection, leading to downregulation of critical genes involved in host defense. These effects were more pronounced in hNECs from females, both by magnitude of effect and number of affected genes. WSP derived from eucalyptus showed a similar trend but with more modest effects while DEP exposure had little effect on gene expression during SARS-CoV-2 infection. We also found that SARS-CoV-2 infection alone altered gene expression patterns differently in hNECs from males and females, with cells from males initiating a more robust upregulation of host defense genes in response to infection. Together, these data support the notion that inhalation of wildfire smoke could adversely affect the host antiviral response to SARS-CoV-2 infection.

Our data indicate that SARS-CoV-2 induced gene expression changes in hNECs are sex-dependent, alone and in the context of WSP exposure. In response to infection, expression of many of the genes in our panel increased, matching previously reported findings about the cellular responses to SARS-CoV-2 infection. For example, induction of type I and type III interferons is well-documented in the epithelial cell response to SARS-CoV-2 infection ((40, 41) reviewed in (42, 43)). We observed significant upregulation of *IFNB1, IFNL1*, and *IFNL2* by 72 h p.i., while *IFNA1* and *IFNA2* were not induced. Accordingly, several interferon-stimulated genes ((44), and reviewed in (45)), regulatory factors, and related transcription factors were also induced in infected hNECs from one or both sexes, such as *ACE2, IFIT1, IFITM3, MX1, DDX58, IRF1, IRF7, STAT1*, and *STAT2*. In addition to activating the interferon response pathway, SARS-CoV-2 is known to activate NF-κB transcription factors and result in upregulation of cytokines and chemokines to recruit immune cells, such as *CSF2, IL6, IL1B, TNF, CXCL8, CXCL9, CXCL10, CXCL11, CCL2, CCL3*, and *CCL5* (41, 46, 47) and reviewed in (48–51). While in our model SARS-CoV-2 infection induced expression of many of these cytokines in both sexes, by 72 h p.i. hNECs from males displayed upregulation of antiviral and immune signaling gene expression which was two times greater than gene induction in hNECs from females, on average. In contrast, in our previous study examining nasal mucosal immune responses to inoculation with live attenuated influenza virus (LAIV) vaccine, Rebuli, et al. observed a more robust antiviral and inflammatory response in female subjects exposed to LAIV compared to male subjects (31). In that study, it was hypothesized that the seemingly larger upregulation of genes involved with antiviral defense and immune cell recruitment in females could reflect differential baseline gene expression levels between the sexes (31). However, in the data presented here, no differences in baseline gene expression between the sexes were observed at 24 and 72 h p.i. This previous *in vivo* human study also revealed that exposure to woodsmoke (500 μg/m^3^) for 2 h prior to inoculation with LAIV resulted in upregulation of inflammatory gene expression in males and suppression of antiviral defense genes in females (31). The data presented here showed a similar, sex-dependent response to woodsmoke exposure in the context of infection. Downregulation of genes involved in the interferon response pathway was more frequent and greater in magnitude in hNECs from females versus males treated with WSP before SARS-CoV-2 infection. Signaling molecules involved in recruitment of immune cells were also generally more downregulated in hNECs from females compared to males. These findings suggest that WSP exposure may dampen antiviral responses in females. Risk assessment studies have found that women more than men are exposed to high levels of PM from burning biomass in indoor settings, especially in developing countries (52–54). However, the opposite is true for exposure to wildfire smoke in firefighters, who are predominantly male. Furthermore, since many of the genes assayed in this study are involved in general antiviral host defense, these results may translate to other viral pathogens of public health importance and additional studies examining sex-based differences in epithelial responses to respiratory viruses alone or in the context of ambient pollutant exposures are needed.

While exposure to WSP significantly modified SARS-CoV-2 induced antiviral host gene expression, the exposures alone in the absence of infection had only modest effects. Exposing hNECs to WSP had few effects on the expression of genes in our panel, besides upregulation of pro-inflammatory *IL6* and *IL1B* and downregulation of *IFNG*. This was not surprising, however, because *in vitro* studies with epithelial cells and *in vivo* human studies have provided inconclusive evidence about the effects of WSP exposure on the airways in the absence of a secondary stimulus. Some studies of human volunteers exposed to WSP did not show significant pro-inflammatory changes in the airway (55–57) while others found signs of pulmonary or systemic inflammation post WSP exposure (11, 58). Pro-inflammatory effects of WSP on epithelial cells *in vitro* are mild and inconsistent (59–61). These discrepancies could be due in part to differences in fuel types and burn conditions across studies. Kim, et al. reported that both burn fuel and temperature affected chemical composition and thus toxicity of WSP in an *in vivo* mouse exposure (7). In our study, WSP from eucalyptus and red oak demonstrated differential effects on gene expression in hNECs. WSP derived from burning red oak contains higher mass fractions of N-alkanes, inorganic and ionic species compared to WSP from eucalyptus (7). Recent data indicate that N-alkanes in PM exposures were significantly associated with inflammation (62), which is consistent with our observations. Additionally, eucalyptus WSP contain a higher mass fraction of methoxyphenols than red oak WSP, which were negatively correlated to biological response in a separate study (7, 63). Hence, the differences in chemical composition and particle size distribution ([Supplemental Fig. S1](https://figshare.com/s/6ba4f3191158622d0d8e)) could be responsible for the differential effects of WSP from red oak and eucalyptus on hNECs.

Even though our data did not show significant differences in viral titers based on sex or particulate exposure, gene expression correlated significantly with viral titers and uncovered positive and negative associations with immune and inflammatory genes. As expected, correlation between viral titer and expression of SARS-CoV-2 N1 and N2 genes was high, indicating viral replication was contributing to viral load, which increased with duration of infection. In addition, expression levels of several IFNs (*IFNB1, IFNL1, IFNL2*) and ISGs (*IFIT1, IFITM3, ACE2, MX1, STAT1, DDX58, CXCL10*, etc.) were positively correlated with viral titer, which has been previously reported (40, 41). In contrast, *TMPRSS2* expression was negatively correlated with viral titer, which was also shown by Lieberman, et al. (41). Interestingly, *IL1B* expression was negatively correlated with viral titer in our model, and expression of *IL6*, *TNF*, and *CXCL8* showed weak positive or no associations with viral titer (*r* of 0.42, 0.14, and 0.28 respectively). These findings may be indicative of viral evasion of pro-inflammatory cytokine induction. Of the genes encoding surfactant proteins, *SFTPA1* was much more strongly associated with viral titer than *SFTPD* (*r* of 0.69 compared to 0.32), although the proteins encoded by both genes contribute to the host immune response (reviewed by (64)). Similarly, *MUC5AC* was more strongly associated with viral titer than its counterpart *MUC5B* (*r* of 0.53 compared to 0.17). Expression of viral and interferon-related genes was highly correlated with viral titer, however inflammatory gene expression generally showed weak or no association with titer. These data indicate that the gene expression response to SARS-CoV-2 infection in our nasal epithelial model is dominated by the IFN response.

The fact that there were no differences in viral load recovered from exposed and unexposed hNECs, even at 72 h p.i., points at some potential limitations of the data presented here. The first is that the changes observed in gene expression at the transcript level may not translate into functional differences at the tissue level. Although *IFIT1, IFITM3, IFNB1, IFNL1, IFNL2, MX1, CXCL10, DDX58*, and other crucial genes for the antiviral response were all downregulated by particulate treatments (in hNECs from females), further investigation is necessary to determine whether these changes result in host defense decrements *in vivo*. Previously, we found agreement between transcript and protein level changes in gene expression after red oak woodsmoke exposure and LAIV inoculation in men and women (31). Further, while the respiratory epithelium represents the first line of defense to inhaled pollution and pathogens, clearance of infection and inhaled debris relies heavily upon recruitment and activation of immune cells. In our study, particulate treatment prior to infection decreased expression of several important chemokines by 72 h p.i. (Fig. 6). It is possible that *in vivo*, the WSP-induced reduction in expression of *CCL3*, *CCL5*, *CXCL10, CXCL11, CXCL9, IL6*, and *TNF*, all of which are chemoattractants for innate and adaptive immune cells, would result in a more widespread and lasting infection. *In vivo* exposures of mice to diesel exhaust prior to respiratory viral infection increased viral titers or viral mRNA collected from whole lungs (65, 66). Management of viral load mediated by immune cells is not captured in our monoculture model. Finally, many groups have reported effective evasion of interferon and NF-κB pathway activation by SARS-CoV-2 (67–70). Indeed, only a small fraction of infected epithelial cells express the majority of interferons and ISGs (40). This suggests that the virus successfully evades or inhibits antiviral responses in the majority of cells it infects. Additionally, Fig. 3 and Fig. 5 suggest that viral replication and release was underway by 24 h p.i., though ISGs and pro-inflammatory responses were not yet induced. The kinetic delay in cellular responses relative to viral replication as well as antiviral evasion by SARS-CoV-2 likely significantly influence the effects of co-exposure to inhaled pollutant on host responses.

Further work is necessary to elucidate the effects of WSP exposure on SARS-CoV-2 infection, especially in other airway cell types and with varied or mixed fuel sources. Exposure to red oak WSP prior to SARS-CoV-2 infection dampens expression of antiviral and host defense genes in nasal epithelial cells. These effects are sex-dependent, with overall greater downregulation of genes in females than in males. Men have been found to be more susceptible to severe and fatal cases of COVID-19 (18). It is possible that wildfire-derived PM could increase COVID-19 morbidity in exposed females, but additional epidemiological studies are needed. The impact of wildfire smoke on public health in the United States and abroad is expected to increase as wildfire seasons become more intense and the population exposed to wildfire smoke continues to rise (4). As viral pandemics and wildfire exposures continue to be concurrent respiratory health risks, it is important to understand the impacts air pollution from wildfires has on host defense responses so strategies for mitigating risk can be employed, especially for subpopulations already susceptible to respiratory infections.

## Code Availability

SAS and R codes used for data processing, statistical analysis, and data visualization are provided as a [Supplemental File](https://figshare.com/s/6ba4f3191158622d0d8e).

## Acknowledgments

The authors would like to acknowledge and thank the study coordinators Noelle Knight, Carole Robinette, and Martha Almond for recruiting hNEC donors and retrieving scrape biopsies. The authors would like to thank Shaun McCullough for his generous donation of DEP and Eva Vitucci for her help in making the DEP preparation and determining particulate sizes. The authors would also like to thank Yong Ho Kim and Ian Gilmour for their generous contribution of WSP samples. Finally, the authors thank the Advanced Analytics Core for their help and contributions.

## Author Contributions

IJ, SAB, and MTH conceived and designed the study; SAB and STB performed experimentation and collection of data; GTB and SAB analyzed data and prepared figures; SAB, IJ, and NEA conceptualized the manuscript; SAB and IJ drafted the manuscript; SAB, NEA, and IJ provided critical revision of the manuscript and interpretation of findings; SAB, GTB, STB, NEA, MTH, and IJ approved the final version of the manuscript.

